# The *HH-GLI2-CKS1B* network regulates the proliferation-to-maturation transition of human cardiomyocytes

**DOI:** 10.1101/2022.04.05.487243

**Authors:** Christina J. Waldron, Lauren A. Kelly, Yasuhiko Kawakami, Juan E. Abrahante, Alessandro Magli, Brenda M. Ogle, Bhairab N. Singh

**Affiliations:** Department of Biomedical Engineering, University of Minnesota, MN, USA; Departments Genetics, Cell and Developmental Biology, University of Minnesota, MN, USA; University of Minnesota Informatics Institute, University of Minnesota, MN, USA; Department of Medicine, University of Minnesota, MN, USA; Stem Cell Institute, University of Minnesota, MN, USA; Department of Pediatrics, University of Minnesota, MN, USA

**Keywords:** GLI2, Maturation, Proliferation, Cardiomyocytes, CKS1B

## Abstract

Cardiomyocyte (CM) proliferation and maturation are highly linked processes, however, the extent to which these processes are controlled by a single signaling axis is unclear. Here, we find the *Hedgehog (HH)-GLI2-CKS1B* cascade regulates the transition between proliferation and maturation in hiPSC-CMs. Initially, we found a significant enrichment of GLI2-signaling in CMs from patients with ischemic heart failure (HF) or dilated-cardiomyopathy (DCM), indicating initiation of fetal programs in the stressed heart. Developmentally, we showed downregulation of GLI-signaling in adult human CM, adult murine CM, and in late-stage hiPSC-CM. In early-stage, proliferative hiPSC-CM, inhibition of Hh- or GLI-proteins enhanced CM maturation. Mechanistically, we identified CKS1B, a new effector of GLI2 and showed that GLI2 binds the *CKS1B* promoter to regulate its expression. *CKS1B* overexpression in late-stage hiPSC-CMs led to increased proliferation with loss of maturation. Thus, the *Hh-GLI2-CKS1B* axis regulates the proliferation-maturation transition and provides targets to enhance cardiac tissue engineering and regenerative therapies.

## Introduction

Transition of proliferative, immature cardiomyocytes to non-proliferative, mature CMs is critical for the generation of contractile force of the functional myocardium (Ahmed et al., 2020; Karbassi et al., 2020). Proliferation of CMs begins to wane during the first postnatal week concomitant with their maturation (Walsh et al., 2010; Xin et al., 2013). Understanding how CMs transition from proliferation-to-maturation and *vice-versa* is key for successful tissue engineering approaches and regenerative therapies. Induction of CM proliferation via de-differentiation processes or using growth factors including IGF, Wnt, Notch, FGF or cell cycle regulators such as CyclinD1, CyclinD2, CyclinA2 is documented. In addition, the characteristics of mature CMs have been well defined and include sarcomere alignment, rod-shaped cell structure, invagination of transverse tubules, altered ion channel gene expression, and isoform switches of troponin (Kannan and Kwon, 2020; Karbassi et al., 2020; Uosaki et al., 2015). However, the regulatory networks involved in CM maturation are less well defined (Kubin et al., 2011; Mohamed et al., 2018; Zhao et al., 2020). Recently, Serum response factor (SRF), metabolic pathways, electrical stimulation and mechanical stimulation were shown to regulate CM maturation. However, the role of these factors in CM proliferation is yet to be documented (Feyen et al., 2020; Guo et al., 2021; Ronaldson-Bouchard et al., 2018) and whether a distinct set of factors or a top hierarchical network globally regulate transitions between proliferation and maturation is unclear.

Precise control of proliferation and maturation of CMs, derived from induced pluripotent stem cells (hiPSC-CMs), is highly critical as they hold great promise for cardiac tissue engineering, regenerative medicine, and drug screening (Funakoshi et al., 2021; Sadek and Olson, 2020; Yang et al., 2014b). Although significant progress has been made, state-of-the-art protocols yield fetal-like hiPSC-CMs, characterized by their immature phenotype, round shape, sarcomere arrangement, contractility, low sarcoplasmic reticulum (SR) calcium, metabolic activity, and gene expression (Karbassi et al., 2020; Ronaldson-Bouchard et al., 2018; Yang et al., 2014b). Although control of CM proliferation is critical to achieve high cell density in engineered heart tissue and to promote full muscle recovery *in vivo*, maturation of hiPSC-CMs remains central to exploit the diverse potential of iPSC-technology. Recent strategies to induce hiPSC-CM maturation reflect understanding gleaned by developmental biology and include long-term culture, mechanical/electrical stimulation, metabolic media formulations (Feyen et al., 2020; Kamakura et al., 2013; Ronaldson-Bouchard et al., 2018; Zhao et al., 2020) (Y Guo et al., 2020, Karbassi E, 2020, Kacey et al 2019, Yang et al., 2014) and activation of signaling pathways including, thyroid hormone, mTORC1, and several microRNAs, miR-125b-5p, miR-199a-5p, miR-221, miR-222 (Chattergoon et al., 2012; Garbern et al., 2020; Lee et al., 2015). Most studies focused on methodologies to induce CM maturation; very few have considered the transition from proliferation to maturation.

Strategies to induce CM maturation are also critical for heart regeneration, as this involves reversal of mature CM to a proliferative state. While factors to revert adult CM to fetal state are explored to promote CM proliferation (Chen et al., 2021; Kubin et al., 2011), factors to promote their maturation are unexplored. Despite the discovery of several signaling pathways in regulating cardiogenesis, their role in heart regeneration is lacking. For example, the role of hedgehog (HH)-GLI signaling is described in cardiovascular development(Singh et al., 2018; Voronova et al., 2012; Zhang et al., 2001), but its function in postnatal CMs is unclear. Hh-signaling is a receptor-ligand signaling modality, involving Glioma-associated factors (Gli; Gli1/2/3) mediated via binding of Hh ligands with Ptch1/Smoothened receptor complex (Briscoe and Therond, 2013; Hui and Angers, 2011; Singh et al., 2015). Developmentally, global knockout of either *Smo* (*Smo^-/-^*) or *Ptc1 (Ptc1^-/-^)*, or double knockouts of *Shh;Ihh (Shh^-/-^;Ihh^-/-^)* results in embryonic lethality due to defects in heart morphogenesis (Zhang et al., 2001). While Gli1^-/-^ mice show no defects, Gli1^-/-^;Gli2^+/-^ compound mutants mice die immediately after birth and Gli2^-/-^ mice are embryonic lethal by E18.5 (Bai et al., 2002; Park et al., 2000). Similarly, Gli2^−/−^;Gli3^+/−^ compound mutants mice showed defects in cardiac outflow tract (Washington Smoak et al., 2005). Notably, the functions of Gli1 is dependent on Gli2- and/or Gli3-mediated transcription(Ali et al., 2019; Bai et al., 2002). While Gli2 acts as an activator and is a primary mediator of HH signaling, Gli3 acts as a transcriptional repressor in the absence of Hh signaling(Hui and Angers, 2011; Ruiz i Altaba, 1999). In cardiogenic conditions, GLI2 function synergistically with MEF2C to regulate cardiogenesis(Voronova et al., 2012). These studies support the notion that GLI2-mediated HH-signaling is critical for embryonic heart development.

Here, we established the *HH-GLI2-CKS1B* signaling axis as a rheostat used to favor proliferation over maturation and vice versa in hiPSC-CM. Modulation of GLI2 regulatory networks could promote hiPSC-CM maturation via global changes in metabolic and transcriptional regulation. Based on genomics analysis of multistage datasets and failing human heart datasets; we postulate that GLI2-signaling attempts to restore the proliferative environment of the diseased heart. Taken together, our findings support the GLI2-cascade as a potential therapeutic target for cardiac tissue engineering and heart regeneration.

## Results

### Hh-GLI2 signaling is operational during the proliferation-to-maturation transition of postnatal cardiomyocytes

In the neonatal period, CMs are highly proliferative (regenerative window), but this capability diminishes rapidly within one-week of their birth concomitant with their maturation (Kannan and Kwon, 2020; Porrello et al., 2011; Soonpaa et al., 1996). To decipher the mechanism of this transition of the postnatal CM, we first mined the publicly available human CM bulk-RNAseq databases, [deposited to Gene Expression Omnibus (GEO)], obtained from isolated human fetal and adult CM (GEO#GSE156707) (Sim et al., 2021). Initial PCA clustering revealed distinct gene expression signature between human fetal vs adult CM (Figure 1A and Figure S1A). We then applied Gene Concept Network (GCN)-methodology, a more precise algorithm than clustering methodology and performed analysis using the down-regulated and up-regulated transcripts. Based on differential gene regulation analysis, we identified a total of ∼8,840 differentially expressed genes (DEGs); of these ∼93.1% (8,232 genes) of the transcripts were downregulated, whereas; ∼8.6% (608 genes) transcripts were upregulated, in adults human CMs (Figure 1A). GCN-analysis revealed robust suppression of multiple cell cycle nodes (Figure 1B, C and Figure S1B) and enrichment of several interconnected metabolic pathways (Figure S1C). Next, we utilized murine datasets obtained from mouse neonatal [postnatal day 1(P1)] and adult (P56) CMs (GEO#GSE95764) (Quaife-Ryan et al., 2017). Differential gene regulation analysis identified a total of ∼9,080 DEGs; of these ∼75.3% (6,841 genes) of the transcripts were downregulated, whereas; ∼24.6% (2,239 genes) transcripts were upregulated (Figure S1D).

**Figure 1.**
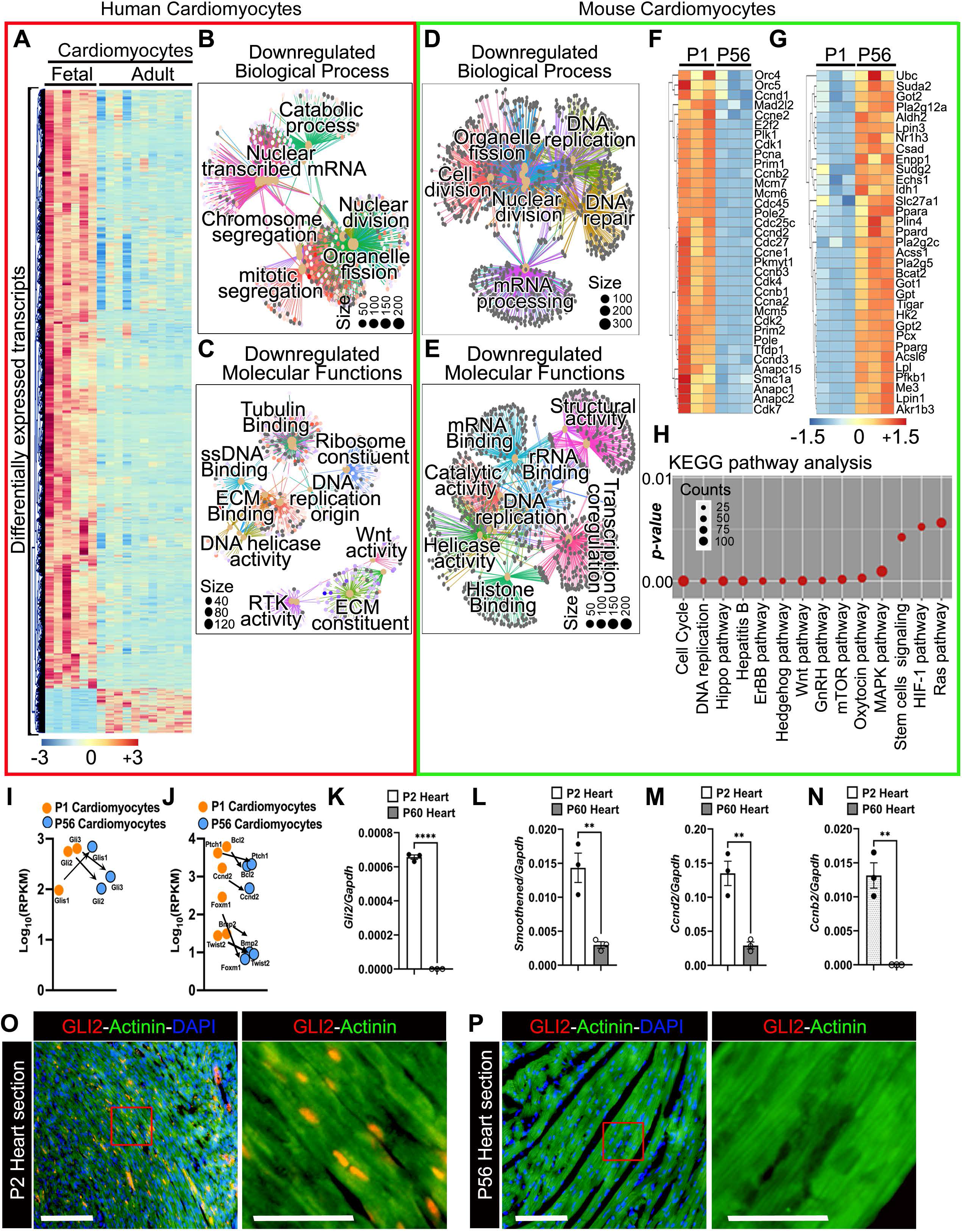
HH-GLI signaling is operational in human and mouse CMs during the regenerative window. **A)** Heat map showing gene expression in isolated human CMs from fetal and adult stage. **(B, C)** Gene-Concept Network (GCN) analysis of significantly enriched biological process and molecular functions in fetal CMs relative to adult CMs. **(D, E)** Gene-Concept Network (GCN) analysis of significantly enriched proliferative and molecular functions nodes in P1 vs P56 CMs. Differential gene expression was determined and adjusted *p-value* cutoff was 0.05 using Fisher’s exact tests. The color in the heatmap indicates the change of normalized gene expression levels. **(F, G)** Heatmaps showing the transcripts up- and down-regulated in P1 and P56 CMs. **(H)** KEGG pathway analysis for highly enriched signaling pathways during the transition from regenerative to non-regenerative window. **(I, J)** Dot-plot showing the expression of GLI-factors and its downstream effectors. **(K-N)** qRT-PCR analysis of *Gli2* and *Smoothened* (*Smo*) and cell cycle regulators, *Ccnd2* and *Ccnb3* from P1 and P60 heart tissue. **(O, P**) Immunostaining using GLI2 and α-Actinin (CMs) antibodies in P2 and P60 mouse heart cryo-sections. The boxed region is magnified in the right panel. Scale bars = 100 μm. Data are shown as mean ± SEM. (n=3 replicates; **p<0.01; ***p<0.0001). (see also Figure S1).

Similar to the human datasets, we found suppression of several nodes related to cell cycle machinery (Figure 1D-F and Figure S1E) with concomitant enrichment of interconnected metabolic cascade (Figure 1G and Figure S1F). This analysis led us to visualize the interactive nodes between networks and their status and showed a global change during the transition from human fetal to adult stage CMs and P1 to P56 stage mouse CMs (Figure 1A-G and Figure S1A-F). To determine the networks operational in fetal/neonatal vs adult CMs, we undertook Kyoto Encyclopedia of Genes and Genomes (KEGG) pathways enrichment analysis for signaling pathways. In agreement with the GCN analysis, KEGG analysis and Gene Ontology (GO)-term analysis showed dysregulation of pathways related to cell cycle and DNA replication in both human and mouse system (Figure 1H and Figure S1B). Among the top 10 annotated signaling pathways, the most significant were ErbB signaling, Hippo signaling, Hh/Gli-pathway, Wnt signaling and mTOR pathways (Figure 1H). While the role of ErBB signaling, Hippo signaling, and Wnt signaling is documented(Sadek and Olson, 2020; van Amerongen and Engel, 2008; Xin et al., 2013; Zhao et al., 2020), the function of Hh/Gli-pathway is not completely defined in postnatal CMs.

To decipher the functions of the GLI-network in postnatal CM, we initially focused on the GLI-members and downstream effectors. Based on the published bulk-RNAseq analysis (Quaife-Ryan et al., 2017), we found high expression (RPKM) of Gli1/2/3 in P1 CMs (Figure 1I) and HH-pathway members (Figure S1G). While the expression of *Gli2* and *Gli3* were reduced, the level of *Gli1* transcripts was increased in P56 CMs (Figure 1I). Similarly, expression of known GLI-mediated downstream effectors including, *Foxm1*, *Bcl2*, *Bmp2*, *Ccnd2* were reduced in P56 CMs relative to P1 CMs (Figure 1J). To validate these omics finding, we performed qPCR analysis using RNA isolated from P2 (regenerative) heart and P56 (non-regenerative) heart. We found robust expression of *Gli2* and *Smoothened (Smo)* with higher levels of cell cycle transcripts including *Ccnd2* and *Ccnb2* in P2 heart (Figure 1K-N) relative to P56 hearts (Figure 1I-L, n = 3 replicates/stage; **p < 0.01; ****p < 0.0001). Next, immunohistochemical analysis using GLI2 antibodies in mouse heart sections revealed expression of GLI2 in the neonatal stage with no detectable expression in the adult heart (Figure 1O, P). These results support that Hh/Gli2-signaling is operational in regenerative/proliferative CMs.

### GLI2-signaling regulates proliferation in postnatal mouse and human cardiomyocytes

To evaluate the functions of Hh signaling in postnatal CMs, we first utilized a small molecule activator, Sonic Hedgehog Agonist (SAG; Hh Agonist) on isolated mouse neonatal CMs and assayed for a proliferative response. Immunostaining for Ki67 together with α-actinin revealed a few CMs stained for Ki67 (∼1-2%; Ki67^+^-Actinin^+^ cells) in the control condition (Figure 2A-C; n = 4 replicates; *p < 0.05). While the percentage of total Ki67^+^ cells did not change significantly (Figure 2A, 2B; n = 4 replicates), we observed an increase in the number of Ki67^+^-Actinin^+^ cells by ∼2-fold in the Hh Agonist condition (Figure 2A, 2C; n = 4 replicates; *p < 0.05)). Next, our EdU-incorporation assay using cultured neonatal CMs showed a ∼2.5-fold increase in α-Actinin^+^-EdU^+^ cells following Hh Agonist treatment compared to the controls without any significant change in total EdU^+^ cells (Figure 2D, 2E and Figure S2A, S2B; n = 4 replicates; **p < 0.01), indicating CM-specific function of HH/GLI-signaling in the regenerative window. Next, we utilized chemical inhibitor, cyclopamine (CyA, Hh Antagonist)(Chen et al., 2002) in these conditions and measured proliferation. Both immunostaining using Ki67/α-actinin antibodies and EdU incorporation assay revealed that inhibition of HH signaling led to significant reduction in proliferative CMs (Figure 2F-J and Figure S2A, S2B; n = 4 replicates; *p < 0.05). Notably, we did not see significant changes in total Ki67^+^ cells as well as total EdU^+^ cells (Figure 2F, 2G, 2I, and Figure S2A, S2B; n = 4 replicates). These results pointed to CM-specific role of Hh/Gli2-signaling in proliferation.

**Figure 2.**
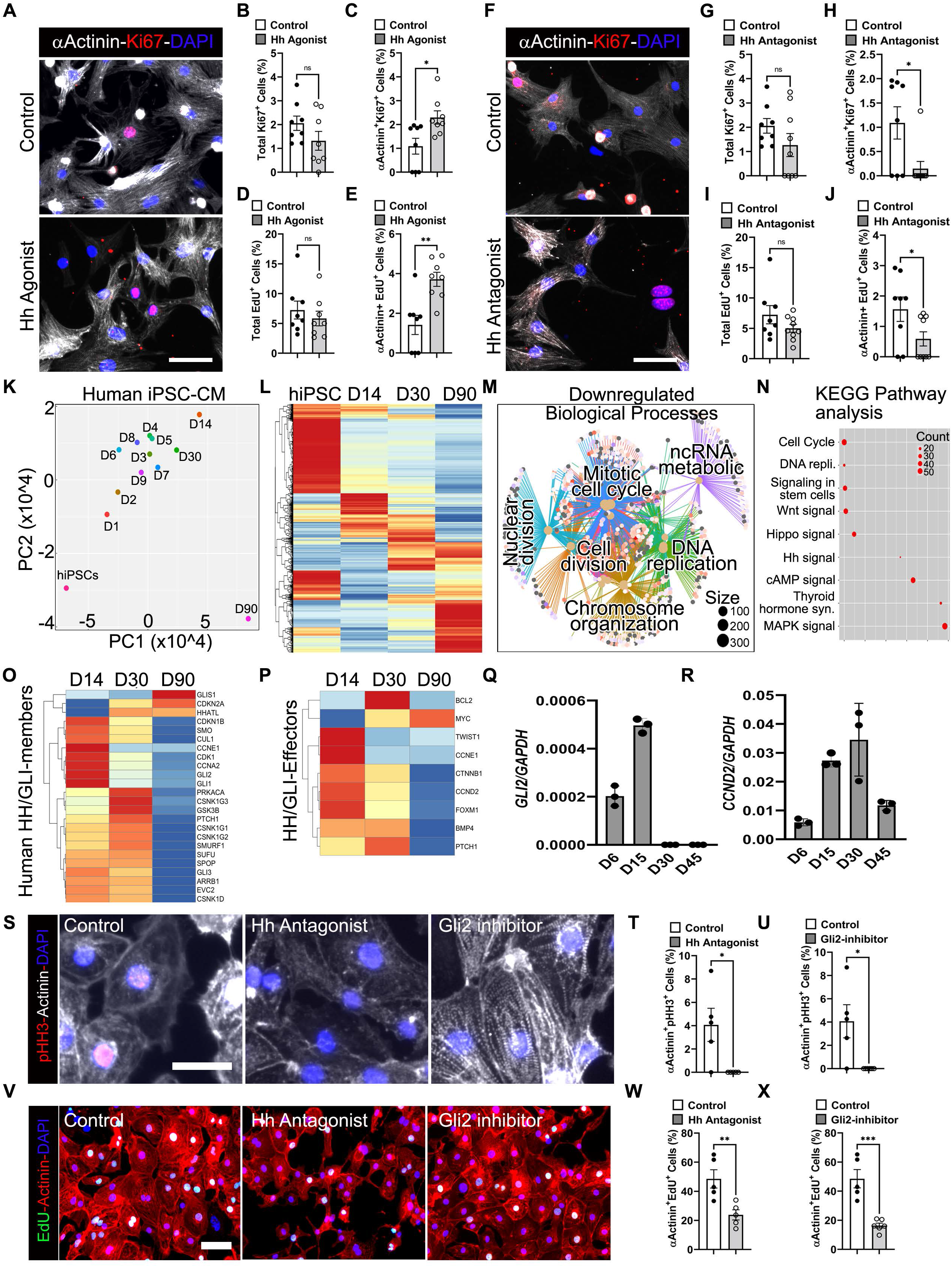
HH/GLI2 signaling modulates proliferation in mouse and human CMs. **(A-E)** Immunostaining using αActinin and Ki67 antibodies (A) and quantification (B-E) of the isolated mouse CMs following treatment with hedgehog (Hh) agonist. **(F-J)** Immunostaining using αActinin and Ki67 antibodies (F) and quantification (G-J) of the isolated mouse CMs following treatment with Hh antagonist (cyclopamine). (n= multiple fields at 20x from 4 replicates). Data are shown as mean ± SEM. **(K)** PCA analysis of bulkRNAseq from hiPSC-CM differentiation between D0-D90 of differentiation. **(L)** Heat map showing the dysregulated gene expression in hiPSC-CM D0, D14, D30 and D90 of differentiation. **(M)** GCN analysis of significantly enriched proliferative networks at D14 relative to D90 hiPSC-CMs. **(N)** KEGG pathway analysis for highly enriched signaling pathways during the transition from immature to mature hiPSC-CMs. **(O, P)** Heatmap showing the expression of Hh and Gli members and downstream effectors at D14 (early), D30 (fetal/neonatal-like) and D90 (late) hiPSC-CMs. **(Q, R)** qRT-PCR analysis of *GLI2* and *CCND2* in differentiating hiPSC-CMs from D6-D45 time points. (n= 3 replicates). Data are shown as mean ± SEM. **(S-U)** Immunostaining using αActinin and pHH3 antibodies (S) and quantification (T, U) of the D20 hiPSC-CM following treatment with DMSO (control), Hh antagonist and GLI2-inhibitor. **(V-X)** Immunostaining using αActinin and EdU staining (V) and quantification (W, X) of the hiPSC-CM following treatment as described in S-U. Data are shown as mean ± SEM. (n= multiple fields at 20x from 5 replicates). Data are shown as mean ± SEM. (ns=non-significant; *p<0.05; **p<0.01). (see also Figure S2).

Next, we sought to examine the potential role of HH/GLI-signaling in hiPSC-CM. Principal component analysis (PCA) of the 13 hiPSC-CM differentiation time points, Day 0 (D0) to D90, revealed distinct clustering of cardiac progenitors that were completely distinct from D0 (hiPSC) and D90 hiPSC-CMs (Figure 2K and Figure S2C). Next, we performed differential expression analysis using 4-stages of hiPSC-CM differentiation, including, D0 (undifferentiated), D14 (early immature CMs), D30 (fetal/neonatal-like CMs) and D90 (late CMs) (Figure 2L). Our analysis revealed unidirectional and gradual transcriptional changes in the gene expression patterns during this time period (Figure S2C). We found a total of ∼12, 590 DEGs; of these ∼52.3% (6,506 genes) of the transcripts were downregulated, whereas; ∼48.6% (6,084 genes) transcripts were upregulated (Figure 2L), in D90 relative to D14 hiPSC-CMS. Further, GCN-based network analysis showed a robust reduction in the cell cycle program at D90 relative to D14 hiPSC-CMs (Figure 2M) with increase maturation programs including ECM dynamics and calcium signaling (Figure S2D). Next, our KEGG analysis revealed a robust downregulation of several signaling pathways including HH/GLI-signals and corresponding downstream effectors (Figure 2N-P and Figure S2E). To confirm these findings, we performed qPCR analysis using RNA isolated from differentiating hiPSC-CMs and found robust expression of GLI2 at early-stage cardiac mesoderm (D6) and in immature CMs (D15) with sharp decline by D30 (Figure 2Q; n = 3 replicates). Similarly, the levels of CCND2 transcripts were high until D30 of hiPSC-CM differentiation with decrease thereafter (Figure 2R; n = 3 replicates). Based on these findings, we predict a critical role of active HH/GLI2-signaling in early hiPSC-CMs proliferation. To evaluate the HH/GLI2-functions in early stage hiPSC-CM (D15-D20), we inhibited HH/GLI-signals using two specific inhibitors affecting distinct levels of the cascade; 1) at the membrane receptor level using CyA; 2) at the GLI2 transcription factor level using Gant61, a specific inhibitor of GLI-proteins (Lauth et al., 2007), and assayed for proliferative indices at D20. Immunostaining using phospho-histone 3 (pHH3; mitosis marker) antibodies together with α-actinin showed ∼4% α-Actinin^+^-pHH3^+^ hiPSC-CMs in the control condition (Figure 2S-U; n = 5 replicates; *p < 0.05). Importantly, we did not detect any α-Actinin^+^-pHH3^+^ hiPSC-CMs following treatment with either inhibitor (Figure 2S-U; n = 5 replicates; *p < 0.05). Next, our EdU-incorporation assay indicated that relative to the control, inhibition using antagonist or GLI2 inhibitor led to ∼2-3-fold decrease in α-Actinin^+^-EdU^+^ cells (Figure 2U-W; n = 5 replicates; **p<0.01; ***p < 0.001). Quantification of α-actinin^+^ cells showed reduced number of hiPSC-CMs following treatment with CyA or Gant61 (Figure S2F, G; n = 5 replicates). These results support a critical role for HH/GLI2 signaling in the regulation of hiPSC-CM proliferation.

### Hh/GLI2-signaling axis regulates cardiomyocyte maturation

To decipher the mechanism by which Hh/Gli2 signals regulate proliferative processes, we performed bulk RNAseq using neonatal mouse CMs following treatment with DMSO (Control), and two distinct small molecule inhibitors of HH/GLI2 signaling 1) HH Antagonist; 2) GLI2-Inhibitor for a 48h period. Relative to the control CMs, we identified ∼13,073 differentially expressed genes (DEGs) with a fold change (FC)>1.2; of these 38% were upregulated genes and 62% were downregulated genes in both the conditions (Figure 3A-D). Compared to Hh antagonist condition, ∼2,500 more transcripts were down-regulated with Gli2-inhibition, whereas, the number of upregulated transcripts were similar in both conditions (Figure 3A, 3C). To identify direct and robust functions of Hh/Gli2 in postnatal CMs, we utilized the common (shared) dysregulated transcripts from these conditions and employed GCN analysis. Our data showed robust downregulation of several nodes related to proliferation processes including, mitotic cycle, DNA replication, and cell division (Figure 3B), further supporting GLI2-signaling during the proliferative phase of CM development.

**Figure 3.**
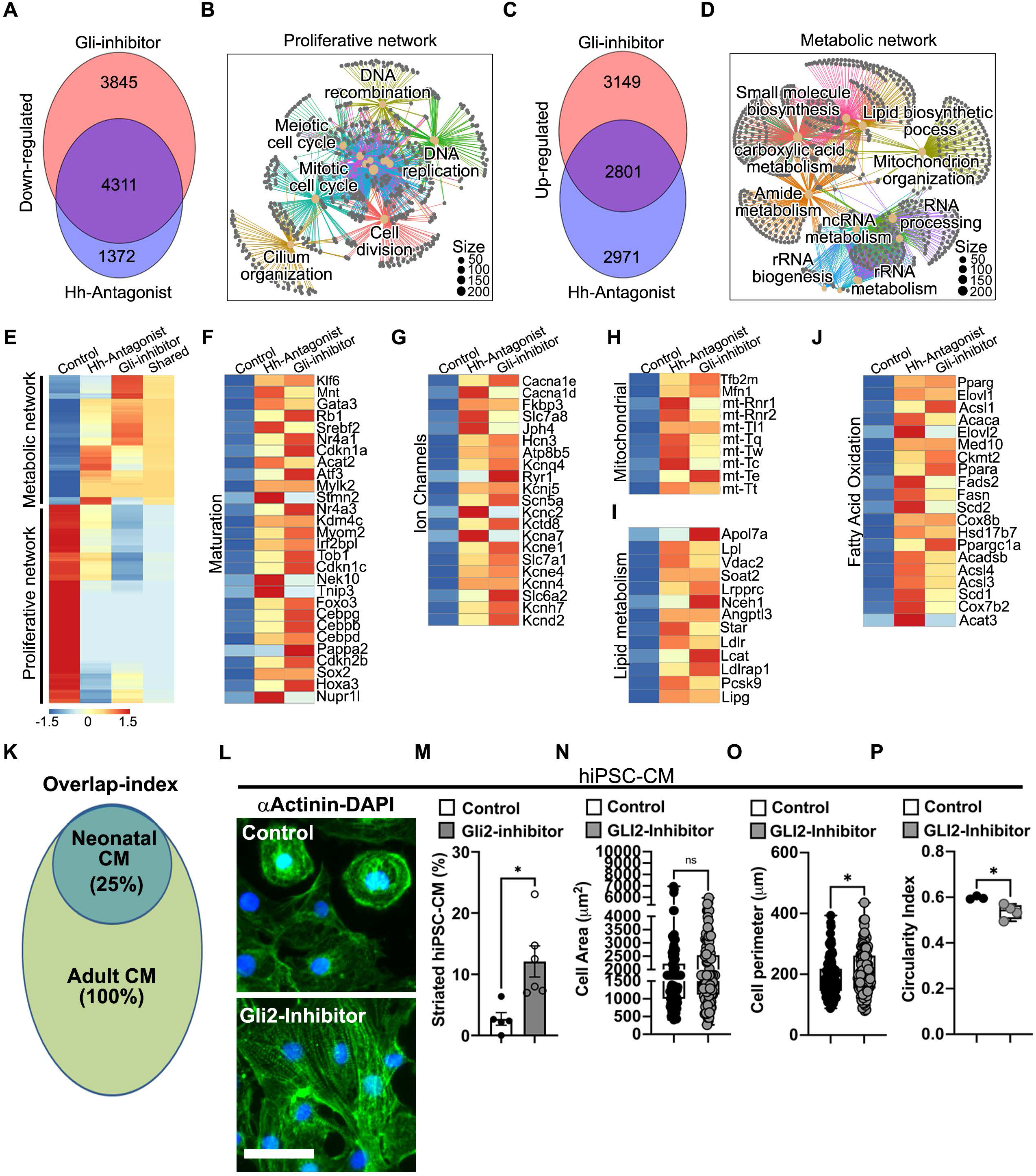
HH/GLI2-signaling axis regulates CM maturation network. **(A)** Bulk RNASeq analysis showing the down-regulated transcripts following treatment with HH antagonist and GLI inhibition. **(B)** GCN analysis of the down-regulated pathways showing reduction in the cell-cycle network following HH/GLI2 network inhibition. **(C)** Bulk RNASeq analysis showing the up-regulated transcripts following treatment with HH antagonist and GLI-inhibition. **(D)** GCN analysis of the up-regulated pathways showing induction of metabolic cascade following HH/GLI2 inhibition. **(E)** Heat map showing the significantly dysregulated transcripts related to metabolic and proliferative network following treatment with HH antagonist and GLI-inhibition. **(F-J)** Heatmap showing the enrichment of maturation programs for selected transcripts for maturation (F), ion channels (G), mitochondrial dynamics (H), lipid metabolism (I) and fatty acid metabolism (J) following treatment with HH antagonist and GLI-inhibition. **(K)** Venn-diagram showing the overlap index of the maturation transcripts induced following GLI-signaling inhibition relative to adult CMs. **(L, M)** Immunostaining using αactinin antibodies (L) and quantification (M) of the striated hiPSC-CMs following GLI2-inhibition. **(M-O)** Quantification of maturation indices following inhibition of GLI2 in hiPSC-CMs; (count n>200 CMs for each condition). Scale bar: 200µm. Data are shown as mean ± SEM. (n = 5 replicates: *p<0.05).

Next, analysis of upregulated transcripts revealed enrichment of several oxidative metabolic cascades following Hh/GLI2 inhibition (Figure 3D). Shift in metabolic program towards fatty acid oxidation is known to be associated with CM maturation(Gaspar et al., 2014). Detailed analysis showed enrichment of transcripts related to CM maturation (Myom2, Mylk2, Rb1, Gata3) (Figure 3E, F), ion channels (Cacna1e, Jph4, Kcnj5, Slc6a2) (Figure 3G), and mitochondrial transcripts (Mfn1, mt-Tl1, mt-Tt, Tfb2m) (Figure 3H). These data indicate an induction of CM maturation following Gli2-pathway inhibition. To further validate the maturation indices, we investigated the carbon-metabolic pathway in CyA/Gant61 conditions as maturing CMs are known to switch from glycolytic to oxidative phosphorylation to meet the increasing energy demands(Guo and Pu, 2020). We found enrichment of several transcripts related to lipid metabolism (Apol7a, Lpl, Soat2, Nceh1) (Figure 3I) and fatty acid oxidation (Pparg, Elovl, Acsl, Med10, Ckmt2) (Figure 3J) following inhibition of the Hh-GLI2 pathway. To visualize the extent of maturation, we performed transcript overlap and found ∼25% overlapping transcripts expressed in the adult mouse CM following GLI2-pathway inhibition (Figure 3K). Developmentally, these changes are associated with CM maturation, therefore, we reasoned that down-regulation of Hh/GLI2 signals could induce hiPSC-CM maturation(Guo and Pu, 2020). To test whether inhibition GLI-signaling could induce maturation in early-stage hiPSC-CMs, we inhibited GLI2-cascade in the immature hiPSC-CMs (D20) and assayed for maturation hallmarks. Our analysis showed increased number of striated hiPSC-CMs, indicating an induction of maturation program (Figure 3L, M; n = 5 replicates; *p<0.05). While the hiPSC-CM area was not changed, we found significant increase in the cell perimeter (hypertrophy) and reduction in the circularity index (higher rod-like CMs) following GLI2 inhibition relative to control (Figure 3N-P; n = 5 replicates; *p<0.05). Overall, these data indicated that GLI2-proteins could regulate the maturation of hiPSC-CMs by globally modulating the transcriptome.

### GLI2-CKS1B cascade regulates the cardiomyocyte proliferation-to-maturation transition

To identify novel downstream effectors and elucidate the mechanism of Gli2-mediated regulation of the proliferation-to-maturation transition, we first utilized the GLI2-ChIPseq datasets obtained from two distinct stages of mouse development (Coquenlorge et al., 2019; Muthu et al., 2019). Our analysis revealed ∼3,977 at E10.5 vs ∼13,012 at E17.5 (Figure S3A).To identify bona fide downstream targets, we explored the common targets from these stages and identified ∼969 targets with Gli2 ChIPseq peaks (Figure 4A). We verified the ChIPseq peaks by exploring the Ptch1 locus (known GLI2 target) and found a robust peak in its regulatory region (Figure S3B). Next, GO-term analysis using the common targets revealed processes required for cardiovascular development, cell cycle, and stem cell maintenance, establishing the role of the GLI-cascade in cardiac development and proliferation (Figure S3C). Next, to visualize the distribution of the peaks, we utilized PAVIS software and found GLI2-ChIPseq peaks in i) ∼37% upstream regulatory regions; ii) ∼19% downstream regions; iii) ∼4.3% in the exons; iv) 0.4% 3’ UTR; and v) ∼10.2% 5’UTR regions (Figure 4A). These analyses showed that Gli2 could bind to multiple regions of the genomic locus and regulate associated expression and functions. We then established the following criteria to identify novel Gli2-targets in CMs: i) high expression in the CMs, ii) presence of Gli2-ChIPseq peak in the upstream regulatory regions, iii) presence of conserved binding Gli2-motif, and iv) target annotated for cell cycle regulation. Next, we employed a 2^nd^ level of filtering using clustering analysis of the down-regulated transcripts from our bulk-RNAseq data using two criteria, i) >1.2-fold downregulation in both HH antagonist and GLI-inhibitor condition relative to control and ii) central regulator of the cell cycle machinery. Based on the above-described criteria, we identified CDC28 Protein Kinase Regulatory Subunit 1B (*Cks1b*), as a top ranked candidate with robust GLI2-ChIPseq peak at its promoter region (Figure 4B, C). Functionally, Cks1b forms a complex with SKP1-SKP2-CUL1 as well as catalytic subunit of the cyclin dependent kinases (CDKs) to regulate cell cycle progression(Martinsson-Ahlzen et al., 2008). Knockout of *Cks1b* (*Cks1b^-/-^*) results in growth retardation, and reduced proliferation(Keller et al., 2007). Our qPCR analysis showed that the expression of *Cks1b* paralleled *Gli2* expression with high expression in the P2 heart and diminished expression by P60 (Figure 4D). Next, qPCR analysis using isolated CMs showed high expression of *Cks1b* in P2 CMs and low expression in P60 CMs (Figure 4D), supporting its role in the regenerative window. To monitor whether GLI2 could regulate the expression of Cks1b, we analyzed the upstream region of the Cks1b locus and identified evolutionary conserved GLI2 binding motifs among various species (Figure 4F). These analyses suggested that GLI2 could directly bind to the Cks1b promoter. To verify these analyses, we isolated neonatal mouse CM and performed qPCR for Cks1b transcript following GLI-inhibition. Our analysis showed that inhibition of GLI2 led to downregulation of *Cks1b* expression (Figure 4G). Next, we utilized hiPSC-CM and performed qPCR for *CKS1B* following GLI-inhibition. Similar to the mouse data, we found robust reduction of the *CKS1B* transcripts in these conditions (Figure 4H). These data indicated that GLI2 is an upstream regulator of *CKS1B* in proliferative CMs.

**Figure 4.**
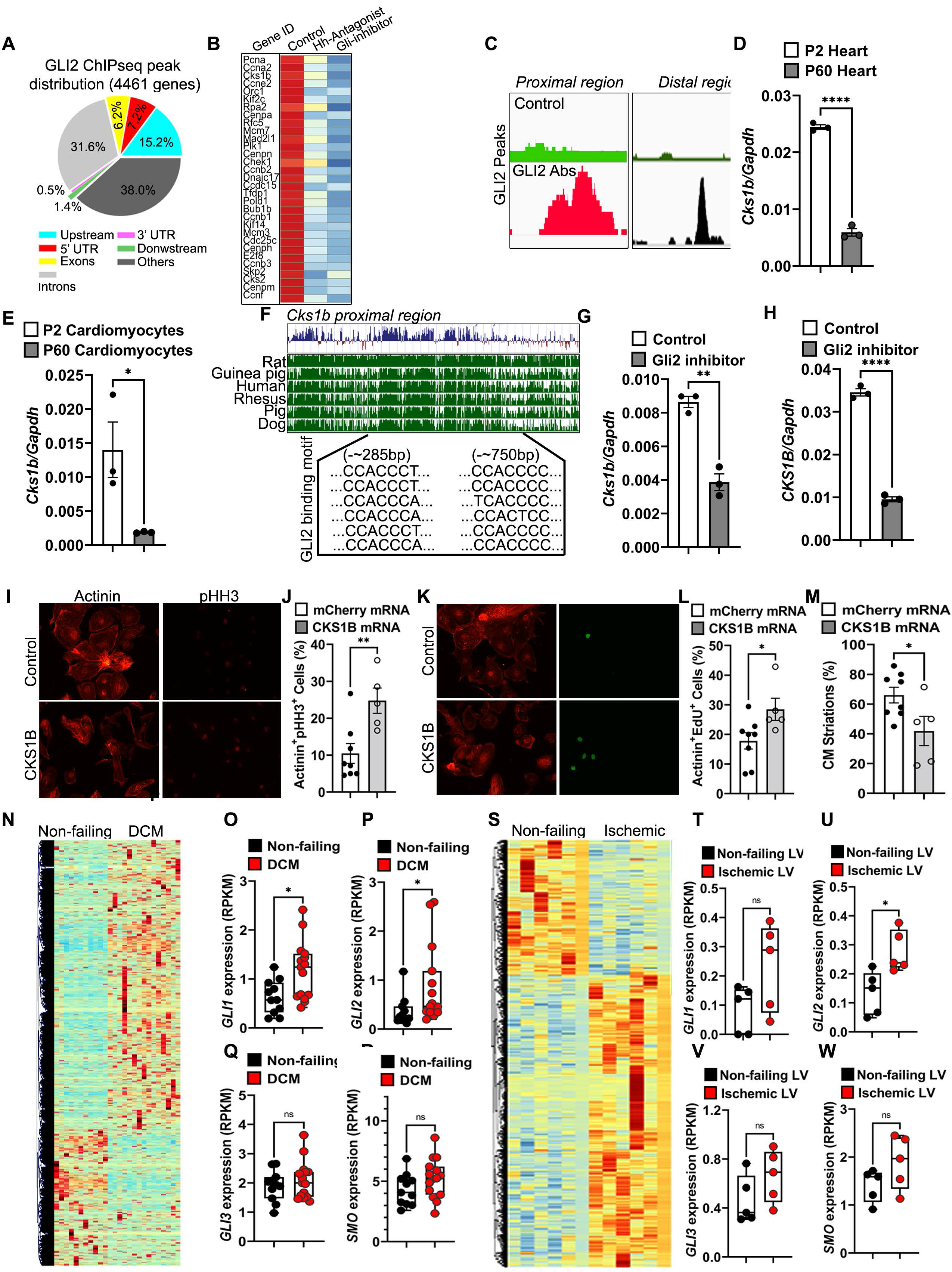
GLI2 is an upstream regulator of CKS1B and GLI2-CKS1B cascade and regulates the CM proliferation-maturation transition. **(A)** GLI2-ChIPseq analysis and Venn-diagram showing the distribution of the Gli2 ChIPseq peaks relative to the TSS using the common targets to visualize the binding of the GLI2 in their genomic regions using PAVIS software. The percent of targets with GLI2-peaks within the various regions of the genome is also shown. **(B)** Heat map showing the significantly dysregulated cell cycle transcripts following treatment with HH antagonist and GLI-inhibition. **(C)** Genomic locus of the Cks1b promoter showing robust GLI2 ChIPseq peaks. **(D, E)** qPCR analysis of Cks1b using RNA isolated from P2 and P60 heart tissue and isolated CMs respectively. Data are shown as mean ± SEM. (n= 3 replicates). **(F)** Evolutionary conservation analysis of the Cks1b promoter region showing the conserved binding sites for GLI2. **(G, H)** qPCR analysis of Cks1b transcripts following treatment with GLI-inhibitor in mouse and human CMs respectively. Data are shown as mean ± SEM. (n= 3 replicates). **(I, J)** Immunostaining using αActinin and pHH3 antibodies (I) and quantification (J) of the hiPSC-CM following 48h post-transfection with control modmRNA and *CKS1B* modmRNA. Data are shown as mean ± SEM. (n= 3 replicates: *p<0.01). **(K, L)** Immunostaining using αActinin and EdU staining (K) and quantification (L) of the hiPSC-CM following 48h post-transfection with control modmRNA and CKS1B modmRNA. Data are shown as mean ± SEM. (n= 3 replicates: *p<0.01). **(M)** Quantification of striated CMs following overexpression of CKS1B relative to control hiPSC-CMs. Data are shown as mean ± SEM. (n= 3 replicates: *p<0.01). **(N)** Heat map showing differentially expressed transcripts from non-failing and DCM heart**. (O-R)** Box plot showing the expression analysis of GLI1, GLI2, GLI3 and SMO in DCM hearts relative to non-failing hearts. (n=11-15 hearts/group; *p < 0.05, ns=non-significant). **(S)** Heat map showing differentially expressed transcripts from non-failing and ischemic heart. (T-W) Box plot showing the expression analysis of GLI1, GLI2, GLI3 and SMO in failing heart relative to non-failing hearts. (n=5 hearts/group; *p < 0.05, ns=non-significant). (see also Figure S3).

Several reports indicated that hiPSC-CMs have limited proliferative capacity with relatively high expression of mature markers by D30 of differentiation. Our analysis showed that both GLI2 and cell cycle transcripts were markedly downregulated by D30 (Figure 2). To evaluate whether the GLI2-CKS1B axis could stimulate proliferation in maturing hiPSC-CM (D30), we synthesized *mCherry* and *CKS1B* modified *mRNA* and perform mRNA transfection experiments followed by a proliferation assay. We observed ∼20% mCherry+ hiPSC-CM at 48h post-transfection (Figure S3D). First, we performed immunostaining using pHH3 (mitotic marker) and α-actinin antibodies and found that mRNA-mediated over-expression of *CKS1B* resulted in a significant increase in the ratio of αActinin^+^pHH3^+^ CMs relative to the *mCherry* transfected cells (Figure 4I, J and Figure S3E; n = 3 replicates; **p < 0.01). Next, we undertook EdU-incorporation assay and performed EdU staining together with α-actinin antibodies. Our data showed a few actinin^+^EdU^+^ cells in the *mCherry* transfected cells (Figure 4K, L). Notably, CKS1B overexpression led to ∼2-fold increase in α-Actinin^+^-EdU^+^ cells (Figure 4K, L and Figure S3F; n = 3 replicates; *p < 0.05)). *CKS1B* transfections led to a significant increase in the number of multi-nucleated CMs (Figure S3G; n = 3 replicates; *p < 0.05). Next, to decipher process by which over-expression of *CKS1B* induced proliferation in mature CMs, we evaluated the sarcomeric status in these conditions. We found that the percentage of striated hiPSC-CM was markedly reduced in the *CKS1B* transfection condition relative to control (Figure 4M; n = 3 replicates; *p < 0.05). These results supports that CKS1B mediates hiPSC-CM proliferation *and* that the GLI2-CKS1B network modulates the proliferative-to-maturation or maturation-to-proliferation transition in hiPSC-CMs.

### GLI2 cascade is re-expressed in human heart patients

It has been shown that both acute and chronic heart diseases result in activation of fetal gene programs (Kubin et al., 2011). Dedifferentiation i.e., loss of sarcomere organization has been documented in patients with DCM and in ischemic heart(Kubin et al., 2011). Several studies have shown that CMs with activated fetal program are more tolerant to ischemic stress. To evaluate whether HH/GLI2-cascade is activated in failing human hearts, we utilized a total of 35 bulk-RNAseq databases obtained from normal (non-failing), ischemic heart failure(GEO#GSE46224) (Yang et al., 2014a) and DCM human patients. By PCA analysis, we filtered the closely clustered datasets and used these for analysis. Clustering analysis of the DCM patients revealed ∼12,246 dysregulated genes (Figure 4N) with downregulation of cell adhesion, PI3-AKT signaling, and MAPK signaling, whereas, upregulated pathways were related to calcium handling, thyroid hormone, vascular smooth muscle (VSM) contraction etc. (Figure S3H). Next, we analyzed the ischemic failing heart vs non failing heart and found dysregulation of similar pathways, suggesting that the failing heart initiates a set of common molecular networks (Figure 4S and Figure S3I, S3J). Analysis of the GLI members in DCM patients showed significant expression of both GLI1 and GLI2 transcripts without any change in the GLI3 levels (Figure 4O-R; n=11-15 hearts/group; *p<0.05), whereas, ischemic datasets showed exclusive and significant activation of GLI2, with non-significant change in the GLI1 and GLI3 levels (Figure 4T-W; n=5 hearts/group; *p<0.05). These data suggest that GLI2-signaling is activated in the diseased heart perhaps to attempt to initiate the regenerative program. Notably, GLI-effectors including CCND2, PTCH (previously known), CKS1B (our discovery) were not significantly changed (data not shown). These analyses indicated that additional co-factors are needed to initiate the regenerative program in the diseased hearts. Further, our analyses provide the basis for exploring GLI2-downstream effectors for more effective therapies for failing heart.

## Discussion

Advancing the field of tissue engineering and regenerative medicine is dependent on precise control of proliferation and maturation of hiPSC-CMs(Huang et al., 2018; Ronaldson-Bouchard et al., 2018; Sadek and Olson, 2020). For instance, promoting CM proliferation could enable heart regeneration following injury(Singh et al., 2018), whereas, promoting maturation of hiPSC-CM will provide better *in vitro* models and improved therapeutics(Huang et al., 2018). Till date, CM proliferation and maturation have been considered as sequential processes(Chattergoon et al., 2012; Uosaki et al., 2015; Zhao et al., 2020), however, we serendipitously find that proliferation and maturation are in fact directly linked by at least one signaling axis, *HH-GLI2-CKS1B.* In particular, we establish the developmental correlation of the axis in CM proliferation, show that inhibition of distinct components of the axis promotes cardiac maturation, and demonstrate the activation of the axis in human heart disease conditions.

Using our bulk-RNAseq data and that from other databases, we established the role of HH/GLI2 signaling in the transition of CMs from proliferation to maturation state. Inhibition of GLI2-factor not only enhanced expression of maturation-related transcripts but also led to increased striations, reduced circularity, increased contractility and higher levels of mitochondrial transcripts. Notably, similar to the GLI2-cascade, Hypoxia Inducible Factor 1a (HIF1a) is shown to be active in fetal cardiac development and is required for maintenance of CM proliferative state. Interestingly, the levels of HIF1a is modulated by media composition, i.e. glucose containing media activated HIF1a levels, whereas, fatty-acid containing media led to its reduction. These studies support the notion that CM maturation is dependent on the carbon source. In further support, our data showed that inhibition of GLI2 results in induction of mitochondrial transcript, fatty acid and lipid metabolic pathways. Therefore, we propose that downregulation of the GLI2-cascade globally modulates maturation kinetics and is critical for maturation initiation.

Next, we established CKS1B as a downstream effector of GLI2 in hiPSC-CMs. CDC28 Protein Kinase Regulatory Subunit 1B (*Cks1b*), is a cell cycle regulator and knockout of *Cks1b* (*Cks1b^-/-^*) results in growth retardation, and reduced proliferation(Keller et al., 2007). We found that over-expression of CKS1B led to reversal of the mature state in hiPSC-CM and induction of proliferative events. Another factor, Oncostatin M is shown to revert the mature state of CM to induce proliferation(Kubin et al., 2011). Recently, Chen et al., have shown reversal of mature phenotype using reprogramming factors to induce heart regeneration(Chen et al., 2021). While these studies support induction of CM proliferation by dedifferentiation and reversal to immature state, our study further extend this inter-linked process, as GLI2 inhibition resulted in induction of the mature state in proliferative CMs. Based on these findings, we propose that the Hh-GLI2-CKS1B axis serve as a rheostat between maturation and proliferation in CMs. Since the levels of GLI2 and CKS1B were low in adult CMs, gene therapy and/or mRNA mediated induction of these factors could initiate adult heart regeneration following injury (Figure S4).

We observed significant upregulation of GLI2 levels in two distinct heart disease condition, ischemia and DCM relative to non-failing hearts. As GLI2-mediated signaling is active in the regenerative stage (<P7), we postulate that the stressed heart attempts to re-initiate the regenerative program by activating the GLI2-mediated fetal program. However, despite the activated GLI2, we observed non-significant changes in other GLI-members and associated downstream effectors. It is possible that a minimum threshold level of GLI2 or other co-factors are necessary to activate the downstream targets. Further, it is also possible that GLI2-signaling synergistically functions alongside other signaling regulators such as Hippo or Wnt signaling to activate the regenerative program. Other studies have also documented similar activation of fetal programs to initially protect stressed hearts(Kubin et al., 2011). Therefore, strategies focused on activation of downstream effectors of GLI2 such as Cks1b could provide new avenues to achieve regenerative therapies for heart diseases.

In summary, our studies support the notion that GLI2 is a prime mediator of proliferative responses in the postnatal CM via a new HH-GLI2-CKS1B cascade. Further, we propose that manipulation of GLI2 allows for the transition between a proliferative and maturation phenotype. In addition to regenerative possibilities, our findings also provide critical insight for tissue engineering approaches. Cardiac tissue engineering approaches has been limited due to lack of high cell density, cellular continuity, and incomplete maturation(Huang et al., 2018). Our study may help alleviating such barriers by promoting a proliferative environment for CMs followed by maturation. To date, stem cell proliferation following by differentiation has been used to achieve high density CMs and synchronous beating in a engineered tissue model(Kupfer et al., 2020). It is possible that modulation of GLI2 signaling pathways could provide additional ways to develop complex engineered heart tissue for disease modeling and drug discovery. Overall, our study enhances understanding of molecular networks involved in postnatal development and in disease conditions. Further, successful induction of such pathways holds unique potential for improved cardiac tissue engineering and induction of cardiac regeneration following injury in humans.

## Methods

Detailed experimental procedures can be found in the supplemental data section.

### Tissue harvest, histology, and immunohistochemistry

All animal experiments were conducted in accordance with guidelines approved by the Institutional Animal Care and Use Committee (IACUC) of the University of Minnesota. Mice were housed in a 12-h light/dark cycle in a temperature-controlled room in the Animal facility University of Minnesota. The day pups were born was considered P0 for their age. For histological analysis, neonatal (P1/P2) mice were euthanized and heart was excised, washed in ice-cold PBS twice and fixed in 4% PFA for overnight period at 4C. For adult heart tissue harvest, animals were perfused with saline, followed by 3M KCl perfusion for 10 min. The hearts were excised using sharp scissor, washed with PBS twice and fixed in 4% for 24h at 4C. The fixed hearts were processed in sucrose gradient (5%-30%), followed by embedding and sectioning. Immunohistochemistry was performed on cryosections (10 µm thick) using standard procedures. Briefly, heart cryo-sections were rehydrated, permeabilized and blocked with 10% normal donkey serum (NDS), 0.1% Triton X-100 in PBS at room temperature and incubated overnight at 4°C with primary antibodies: α-actinin (Abcam; 1:300), GLI2 (Novus biologicals; 1:100). Sections were rinsed and incubated with combinations of secondary antibodies (1:400) including anti-mouse Alexa 488, anti-rabbit Alexa 594 (Jackson ImmunoResearch Laboratories) for 1h at room temperature. Nuclei were stained using DAPI and mounted using wet mount (Invitrogen).

### Mouse ventricular cardiomyocyte isolation using Magnetic Cell Sorting

Ventricular CMs were isolated using previously published protocols. Briefly, ventricles were dissected from P1/P2 pups, wash with ice-cold DPBS twice followed by mincing in calcium and bicarbonate-free Hanks-buffer with 20mM HEPES (CBFHH buffer). Tissue digestion was performed using enzyme solution containing 1.75 mg/ml of trypsin (Difco) and 20 µg/ml of DNaseII (Sigma-Aldrich). Digested cells were washed with ice-cold DPBS and CMs were isolated by magnetic negative cell sorting using CD90 antibodies to remove fibroblasts. Using this protocol, we routinely obtained >90% CMs (confirmed using -actinin antibody immunostaining). We plated 70,000 CMs/well in a fibronectin coated 12-well plate. After 12h, the culture medium (DMEM, 10%FBS, Vitamin B12, 1X penicillin-streptomycin) was changed, and cells were subjected to the different treatments (SAG;4µg/ml, CyA;5µg/ml and Gant61;3uM) for 48h and harvested for further analysis. For the EdU incorporation assay, CMs were incubated with 20µM EdU for 48h and fixed using 4% PFA for 10 min at room temperature.

### hiPSC culture and hiPSC-CM Differentiation

hiPSCs were cultured on matrigel-coated plates in mTeSR1 medium plus supplements (1X) for 3-4 days to attain 80-90% confluency. The culture media was changed every day. Differentiation of hiPSC to CMs were performed using previously published protocol using Wnt modulation protocol (Lian et al., 2012). Briefly, hiPSCs cells were harvested using accutase solution and the singularized cells were resuspended in culture media supplemented with 5 μM ROCK inhibitor (Y27632). 1x10^6^ cells/well of 12-well plate were seeded on matrigel-coated plates and expanded until they reached 90% confluence. Cardiac differentiation was initiated (D0) using RPMI-1640/B27 without insulin (RPMI/B27 minus) containing 8 μM GSK3-β inhibitor (CHIR99021) for 24 hours in the 37°C, 5% CO2 incubator. On D1, the medium was replaced with fresh RPMI/B27 minus medium. At D3, fresh and exhausted medium was mixed in 1:1 ratio and supplemented 1X Wnt inhibitor (IWP2; 5 μM) and added to the cells. The media was changed on D5. From D7 onwards, the media was replaced with RPMI1640 supplemented with B27 supplement containing insulin (RPMI1640/B27-plus). Spontaneous twitching was observed by D8-10, and robust continuous contraction occurred by D12. The media was changed every 3 days for maintenance. To enrich for CM population, hiPSC-CMs were cultured in no glucose-sodium lactate media from D12-D14, and thereafter, cells were maintained in RPMI1640/B27-plus medium.

### hiPSC-CM re-plating and small molecule treatment

To replate, hiPSC-CMs were dissociated using 0.5ml of TrypLE (10X) select for 10-15 min at 37°C. Single cell suspension was obtained by resuspending the hiPSC-CMs in RPMI1640/B27 plus 10% knockout serum media and 90,000 cells/well cells were seeded on Matrigel-coated 12-well plates in RPMI/B27-plus medium containing 5μM ROCK inhibitor. hiPSC-CMs were subjected to the different treatments (DMSO, CyA;5µg/ml and Gant61;3uM) for 48h and harvested for further analysis. For the EdU incorporation assay, hiPSC-CMs were incubated with 20µM EdU for 48h and fixed using 4% PFA for 10 min at room temperature.

### Immunostaining and EdU-incorporation assay

Following treatment with various small molecules, neonatal mouse CMs, hiPSC-CMs were washed with PBS followed by fixation using 4% paraformaldehyde (PFA) for 15 min at room temperature. Fixed cells were permeabilized by 0.2% Triton X-100 at room temperature for 15min, washed with (0.01%) PBST 3 times and blocked using 5% normal donkey serum (NDS) in PBST. Subsequently, cells were incubated overnight at 4 °C with primary antibodies: α-actinin (Abcam; 1:300), α-phospho-Histone H3 (Ser10) (Millipore; 1:100), Ki67 (Abcam; 1:200) in the blocking solution in a humified chamber. Cells were then rinsed and incubated with combinations of secondary antibodies (1:400) including Alexa 488, Cy3, and Cy5 (Jackson ImmunoResearch Laboratories). EdU staining was performed using the EdU labeling kit (Life Technologies). Nuclei were stained using with 0.1μg/mL of DAPI dye for 15 min at room temperature. Cells were kept in PBS at 4°C and imaged using Leica fluorescence microscope.

### RNA Isolation and qPCR Analysis

For RNA isolation from whole heart tissue, P1/P2 and P56 hearts were harvested, washed in PBS and snap-frozen in liquid N2, followed by homogenization using tube-pestle. Subsequently, the tissue powder was lysed using 600ul of RLT-lysis budder as per the manufacturer’s protocol (Qiagen). For total RNA isolation from isolated CMs, hiPSC-CMs, cells were washed with sterile PBS and lysed in RLT-lysis buffer for specific time points using the RNeasy kit (Qiagen) RNA isolation was performed as per the protocol, followed by on-column DNA digestion to remove any traces of DNA as per the instructions. For cDNA synthesis, 100-1000 ng of total RNA was used and synthesized using the SuperScript IV VILO kit (Thermo Fisher Scientific) according to the manufacturer’s protocol. Quantitative PCR (qPCR) was performed using gene-specific oligos and the SYBR-green method. The list of primers used in this study is provided in Table S2.

### Bulk-RNAseq and Bioinformatic analysis

The total RNA was isolated as described above. Bulk-RNAseq experiments were performed at the University of Minnesota Genomics Core (UMGC, UMN). We performed pair-end sequencing with an average of >43 million reads per sample. 2 x 150bp FastQ paired-end reads were trimmed using Trimmomatic (v0.33) enabled with the optional-“q” option; 3bp sliding window trimming from 3’-end requiring minimum Q30. Quality control on raw sequence data for each sample was performed with *FastQC*. Read mapping was performed via Hisat2 (v2.1.0) using the *Mus musculus* genome (GRCm38v100) as reference. Gene quantification was done via Feature Counts for raw read counts. Differentially expressed genes were identified using the Proportional Estimation test based on Kal’s test feature in CLCGWB (Qiagen, Redwood City, CA) using the raw read counts for our bulk RNAseq datasets. We filtered the generated list based on a minimum of 1.2X Absolute fold change and Bonferroni corrected, p<0.05. Differential gene expression was performed using the R package limma (Version 3.28.14) for the available databases. Cutoff values of absolute fold-change ranging from >1.2 – 2.0 and false discovery rate < 0.05 were used to select differentially expressed genes between sample group comparisons. Pathway and gene set enrichment analysis were performed using the R package clusterProfiler (Version 3.0.4). KEGG and GO results (p< 0.05) were visualized using the R package DOSE (Version 3.16.0).

### Data availability

All data presented in this study are available in the main text or the supplementary materials. Raw and analyzed RNA-sequencing data generated during this study are available in the Gene Expression Omnibus (GEO) repository (https://www.ncbi.nlm.nih.gov/geo/) and are accessible through GEO series accession number. Previously published datasets used in this study include: GSE81585, GSE95762, GSE46224, and GSE116250.

### Synthesis and Transfection of *CKS1B* and *mCherry* modified mRNAs

PCR products with the T7 promoter site in the 5′ end for human CKS1B (Primers: T7 forward: AATACGACTCACTATAGGGCACCATGCCCAGCTGCACCGCGTC, CKS1B reverse: TTAGCAAGTCCGAGCGTGTTCGAT) and mCherry (Primers: T7 forward: AATACGACTCACTATAGGGCACCATGAGCGGGGGCGAGGAGCTG, mCherry reverse: TTATCTGAGTCCGGACCTGTACAG), coding sequences were amplified from the hiPSC cells and plasmid vector having coding region for mCherry. Subsequently, PCR products were purified and 500 ng of the PCR template was used for the *in vitro* synthesis and capped using Ultra sysnthesis kit (APEXBio) as per the protocol. The final capped transcription reaction was performed at 37 °C for 14 h followed by the poly(A) tailing reaction as per the manufacturer’s instructions. RNA was recovered using the phenol-chloroform protocol. 1.5 μg of the purified RNA was used for the transfection experiment using ViaFect in D30 replated hiPSC-CM. For the EdU incorporation assay, hiPSC-CMs were incubated with 20µM EdU for 48h and fixed using 4% PFA for 10 min at room temperature and processes further. EdU staining was performed using the EdU labeling kit (Life Technologies).

### Statistical analysis

All experiments were repeated at least three times and the values presented are mean ± standard error of the mean (SEM). Statistical significance was determined using the *Student’s t-test*, *Mann-Whitney Test* for Statistical significance was determined using the Student’s t-test and a p-value ≤ 0.05 was considered as a significant change.

## AUTHOR CONTRIBUTIONS

C.J.W, L.K., B.M.O., B.N.S. conceived and designed the research. C.J.W, L.K., and B.N.S. designed and executed the experiments including bioinformatic analysis, cardiac differentiation, functional measurements, immunostaining, imaging and quantitative analysis. C.J.W, L.K. and B.N.S. performed quantitative analysis and summarized the data. J.E.A.L. helped generate raw data for the RNAseq analysis. A.M. helped with bulkRNAseq, ATACseq and ChIPseq analysis and data generation. C.J.W, B.M.O. and B.N.S. discussed the results and generated the figures. C.J.W, L.K., S.G., J.E.A.L., A.M., Y.K., B.M.O. and B.N.S helped write the manuscript. All authors approved the manuscript.

## ACKNOWLEDGMENTS

We acknowledge the UMGC core facility, animal core facility for their services. This research was supported by Regenerative Medicine Minnesota (RMM 091718 DS 007 to B.N.S. and RMM 091620DS008 to B.M.O. and B.N.S.), and the National Heart, Lung, and Blood Institute of the National Institutes of Health (R01 HL137204 to B.M.O.).

---

**Figure S1.**
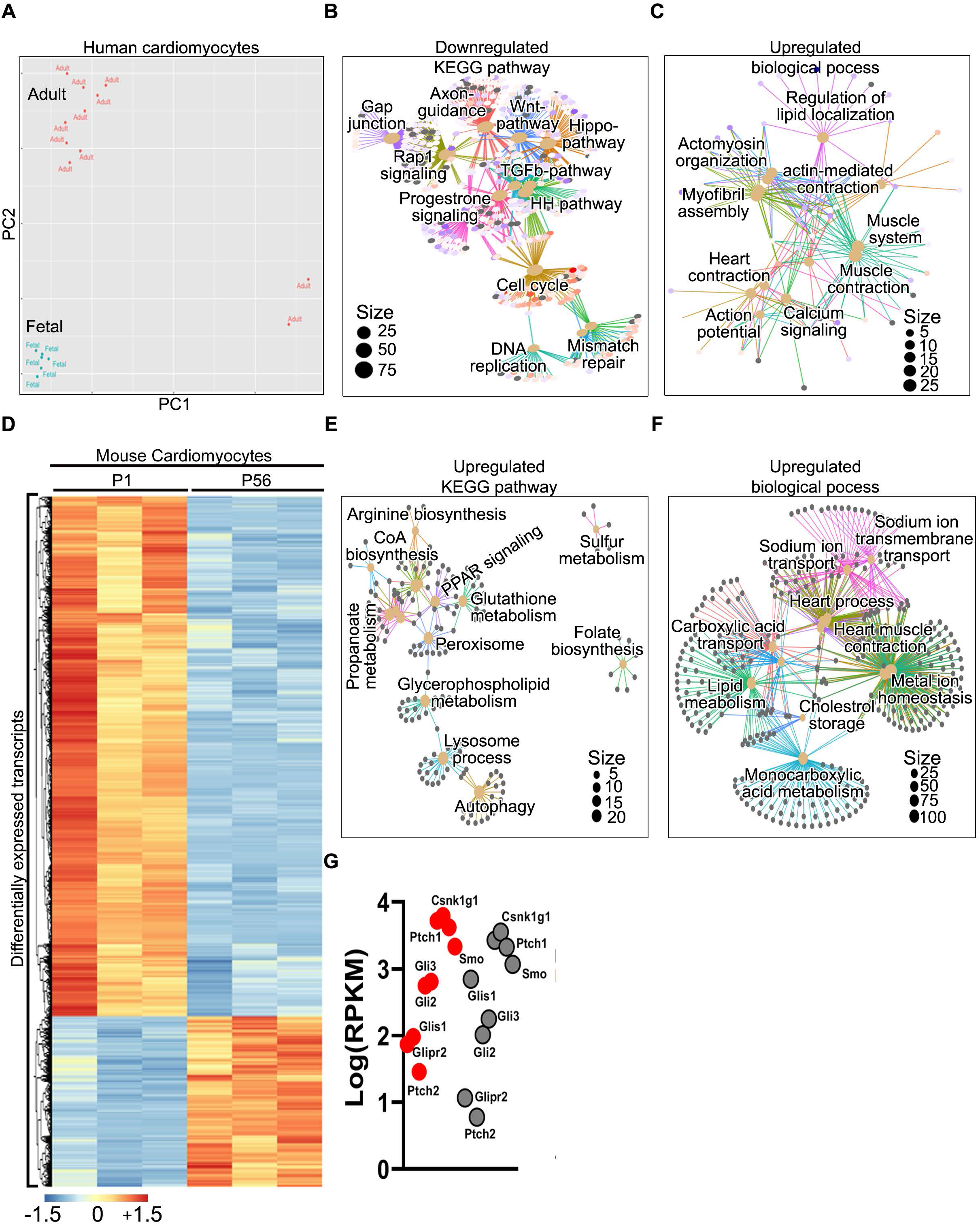
Related to Figure 1. Downregulation of HH-GLI-signaling leads to maturation in postnatal cardiomyocytes. **(A)** PCA analysis of the bulkRNAseq datasets obtained from human fetal and adult CMs. Fetal CMs clustered distinctly from adult CMs, indicating transition from fetal to adult involve different transcriptional profile. **(B, C)** GCN analysis of downregulated KEGG pathways (B) and upregulated biological process (C) in adult CMs relative to fetal stage. Note the close association of cell cycle network with HH pathway (B). **(D)** Heat map showing differentially expressed genes in isolated CMs from postnatal day 1 (P1) vs P56 stage. Note the robust change (up- and down-regulated) in the expression of various genes at these time periods. **(E, F)** GCN analysis of upregulated KEGG pathways (E) and upregulated biological process (F) in adult CMs relative to fetal stage. Note the enrichment of metabolic process in the P56 stage relative to neonatal stage. The dot-size in the GCN analysis indicate the number of associated genes. **(G)** Dot-plot showing the expression values [(Log (RPKM)] of GLI-pathway factors in P1 CMs (red) vs P56 CMs (grey). (n=3 replicates/stage).

**Figure S2.**
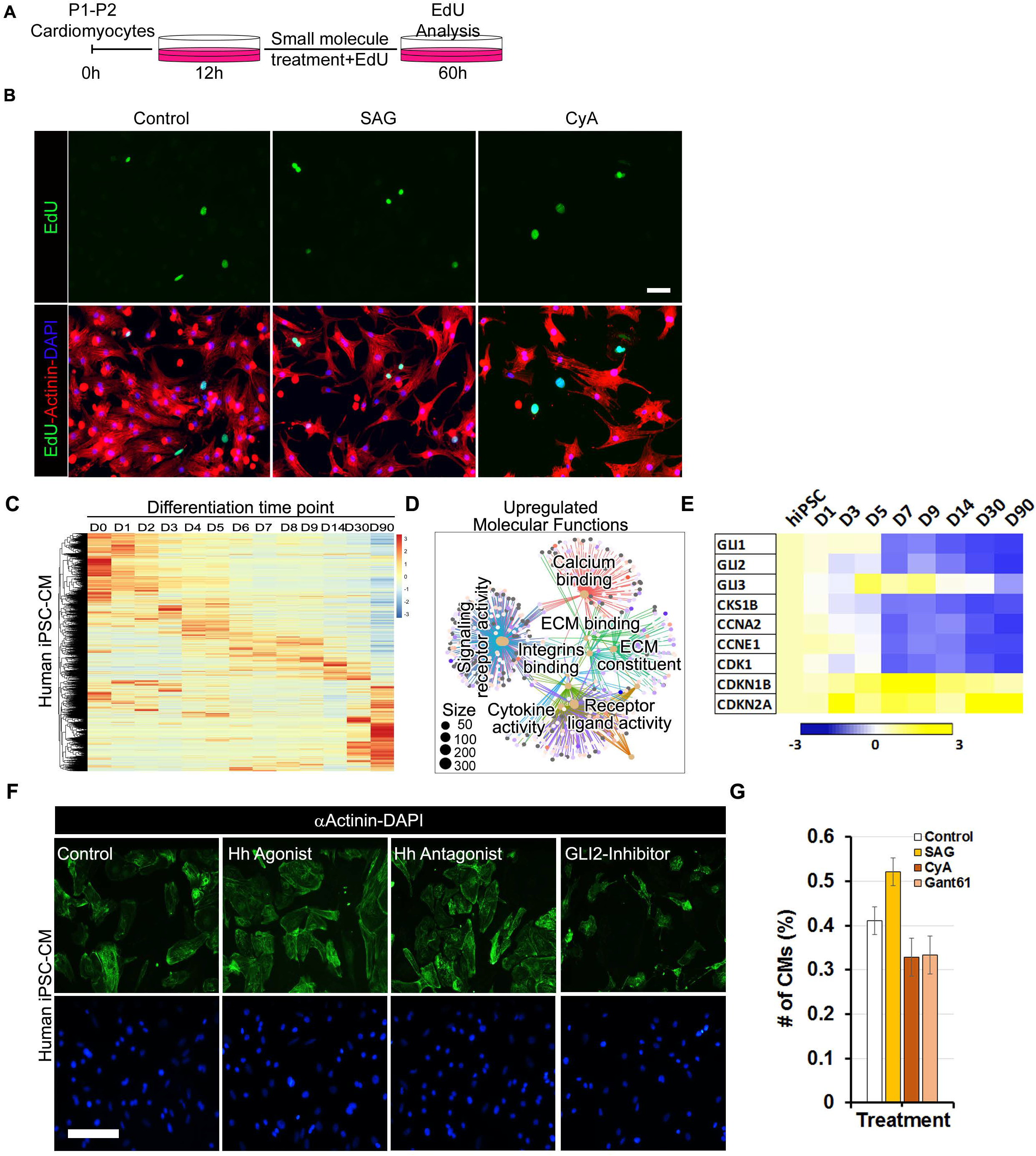
Related to Figure 2. GLI-signaling regulates proliferation in hiPSC-CM at early stage. **A)** Schematic of small molecule treatment for HH/GLI modulation in the neonatal CMs. **(B)** Immunohistochemical images of α-Actinin^+^-EdU^+^ isolated neonatal CMs following exposure to control, SAG or CyA and together with EdU incorporation assay. **(C)** Heat map showing the dysregulated gene expression in hiPSC-CM D0-D90 of differentiation. **(D)** GCN analysis of significantly enriched molecular functions at D90 relative to D14 hiPSC-CMs. Note the increase nodes for ECM proteins and calcium dynamics in the D90 hiPSC-CMs. (E) Heatmap showing the expression of HH/GLI members and cell cycle members between D0-D90 of hiPSC-CM differentiation. Note the downregulation of GLI-members with concomitant increase in cell cycle inhibitors (*CDKN1B, CDKN2A*) at late stage of hiPSC-CMs. **(F, G)** Immunostaining using αActinin antibodies (F) and quantification (G) of the D15-20 hiPSC-CM following treatment with DMSO (control), HH Agonist (SAG), Hh antagonist (CyA) and GLI2-inhibitor (Gant61). Data are shown as mean ± SEM. Nuclei are shown in blue. Scale bar: 200µm.

**Figure S3.**
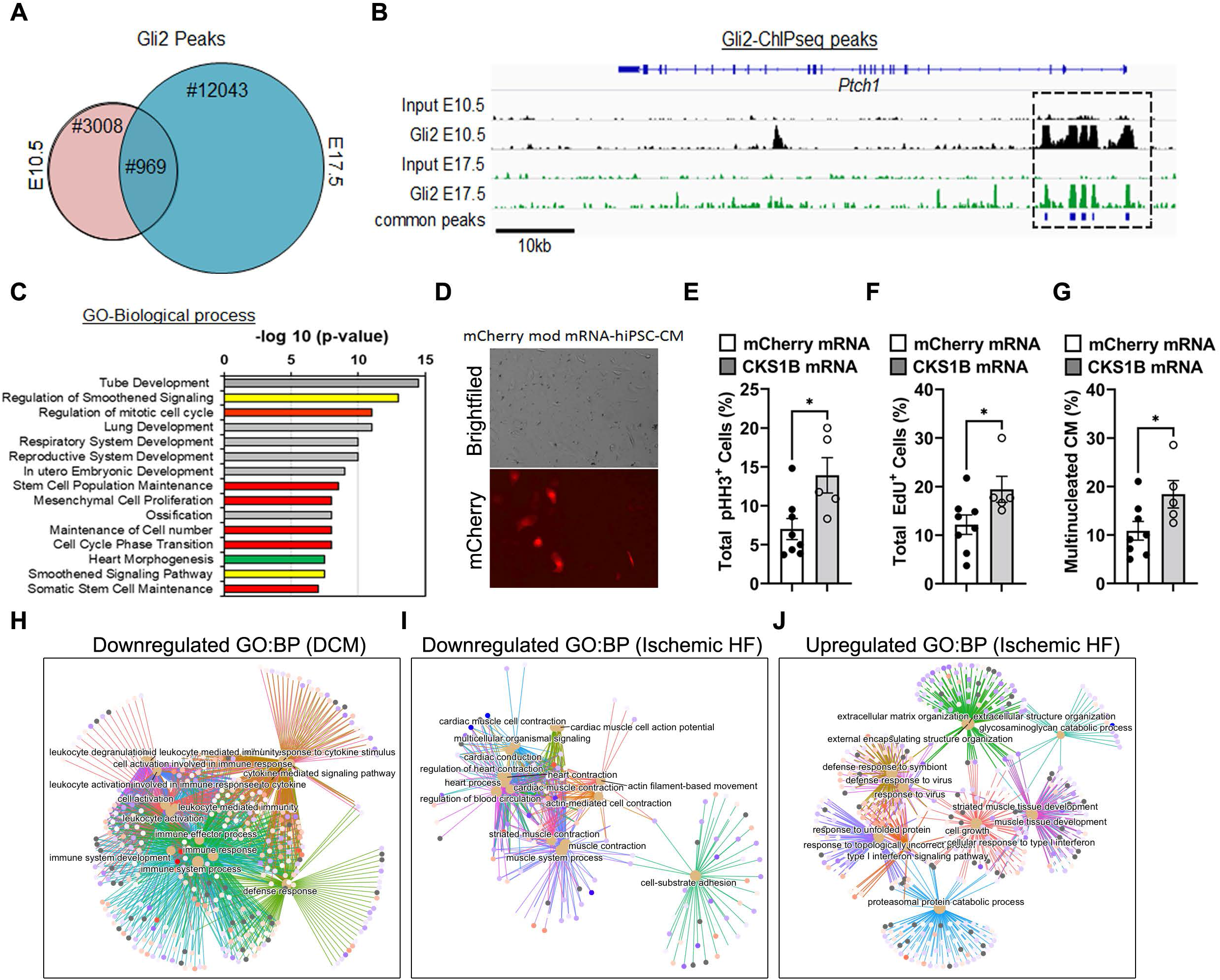
Related to Figure 4. GLI2-CKS1B axis regulate hiPSC-CM proliferation and maturation. **(A)** Venn-diagram showing the Gli2 ChIPseq peaks at E10.5 and E17.5 stages. The intersection shows the common targets from these time points. **(B)** Genomic locus of the *Ptch1* gene with GLI2 ChIPseq peaks at E10.5 and E17.5 time periods. The box region indicates the location of the peaks. **(C)** GO-terminology analysis for biological process using the gene sets obtained from the common targets. Note the enrichment of biological processes involved in smoothened signaling pathway (yellow bars), cell cycle regulatory pathways (red bars) and cardiac morphogenesis pathway (green bar). (D) Brightfield and fluorescence image showing transfection of mCherry modified mRNA 24h post-transfection. **(E, F)** Quantification of total pHH3^+^ (E) and total EdU^+^ (F) hiPSC-CM following 48h post-transfection with control modmRNA and *CKS1B* modmRNA. Data are shown as mean ± SEM. (n= 3 replicates: *p<0.01). **(G)** Quantification of multi-nucleated hiPSC-CM following overexpression of CKS1B relative to control hiPSC-CMs. Data are shown as mean ± SEM. (n= 3 replicates: *p<0.01). **(H)** GCN analysis of the down-regulated biological process in DCM patients relative to non-failing hearts. Note the enrichment of nodes related to immune response and cytokine pathways. **(I, J)** GCN analysis of the down-regulated (I) and upregulated (J) biological process in ischemic failing heart patients relative to non-failing hearts. Note the enrichment of nodes related to cellular adhesion, muscle contraction, actin filament movement were downregulated, whereas, ECM protein organization, IFN signaling, proteasomal pathways were upregulated in the failing hearts.

**Figure S4.**
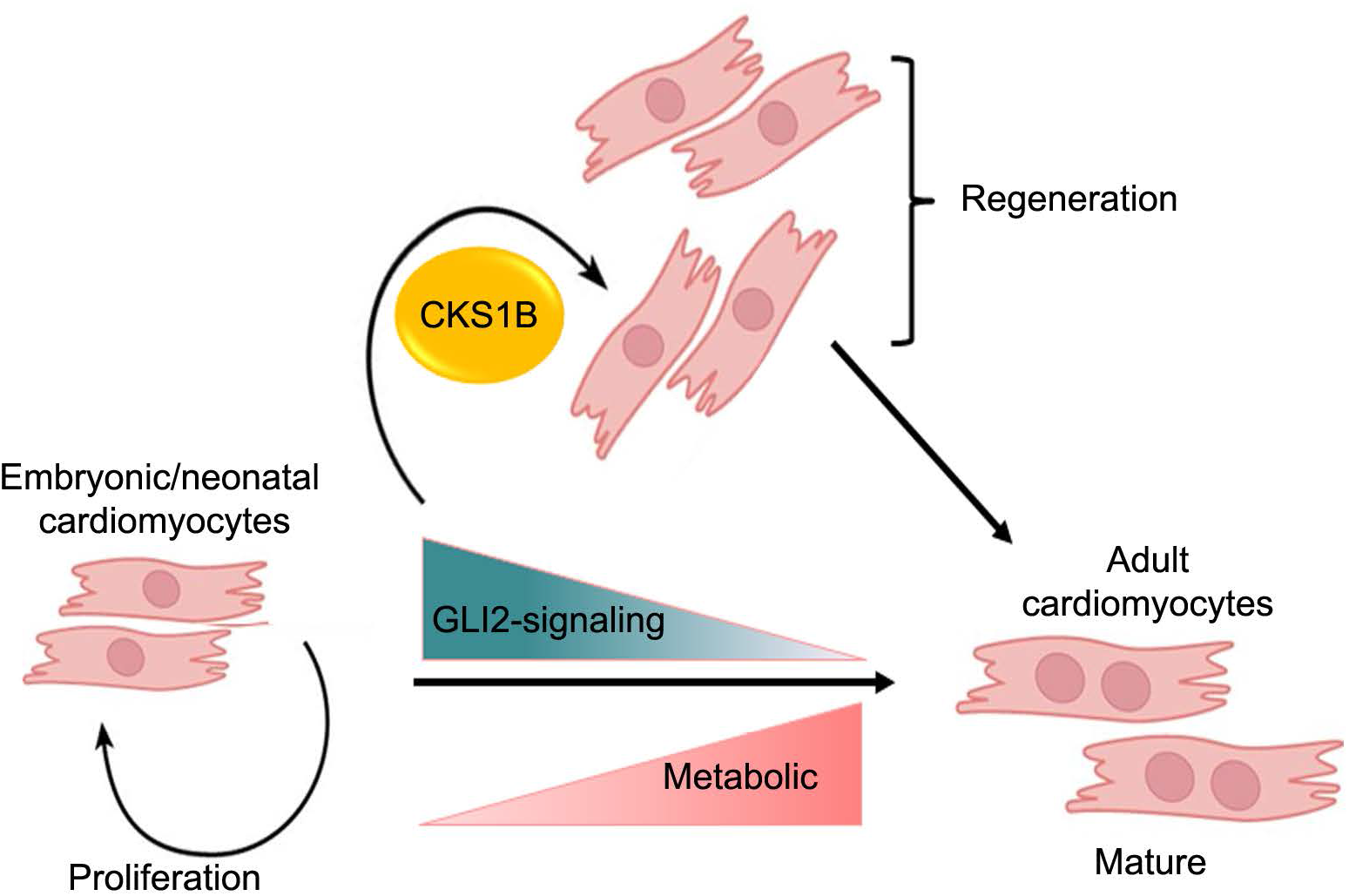
GLI2-CKS1B cascade in proliferation-to-maturation transition. A model summarizing the involvement of HH-GLI2-CKS1B network in the regulation of CM proliferation and maturation transition. GLI2 is primarily expressed in the early neonatal stages and subsequently gets downregulated as the postnatal CM mature with their adaption to metabolic change required to meet the energy demands. Inhibition of GLI2-signaling leads to induction of maturation in the neonatal CMs, whereas, activation of GLI-signaling promotes proliferation via its downstream target, CKS1B, to regulate the proliferative response in CMs. Modulation of the GLI2-CKS1B axis provide new strategy for the transition from proliferation-to-maturation and vice-versa.

